# Hello, is that me you are looking for? A re-examination of the role of the DMN in off-task thought

**DOI:** 10.1101/612465

**Authors:** Charlotte Murphy, Giulia Poerio, Mladen Sormaz, Hao-Ting Wang, Deniz Vatansever, Micah Allen, Daniel S. Margulies, Elizabeth Jefferies, Jonathan Smallwood

## Abstract

Neural activity within the default mode network (DMN) is widely assumed to relate to processing during off-task states, however it remains unclear whether this association emerges from a shared role in self or social cognition. In the current study, we examine the possibility that the role of the DMN in ongoing thought emerges from contributions to specific features of off-task experience such as self-relevant or social content. A group of participants described their experiences while performing a laboratory task over a period of days. In a different session, neural activity was measured while participants performed self/other judgements. Despite the prominence of social and personal content in off-task reports, there was no association with neural activity during off-task trait adjective judgements. Instead, during both self and other judgements we found recruitment of caudal posterior cingulate cortex - a core DMN hub - was above baseline for individuals whose laboratory experiences were characterised as detailed. These data provide little support for a role of the DMN in self or other content in the off-task state and instead suggest a role in how on-going thought is represented.

## 1. Introduction

The human mind is not always occupied by our actions in the moment: everyday experience is replete with self-generated thoughts that are based on representations from memory, rather than events in the here and now (Smallwood & Schooler, 2015). Since the turn of the century, an increasing focus on states of self-generated thought, often under the rubric of mind-wandering (Seli et al., 2018), has revealed links with health, well-being and productivity (Andrews-Hanna, Smallwood, & Spreng, 2014). This research endeavour has been enhanced and supported by the discovery of the default mode network (DMN), a distributed set of regions spanning medial and lateral, frontal and parietal cortices (Raichle, 2015). Unlike other neural systems, the DMN regions deactivate during effortful external processing (Raichle, 2015), and these so-called “task-induced deactivations” were highlighted as a potential analogue to the processes occurring during off-task cognitive states (McKiernan et al., 2003).

Subsequent research broadly corroborated this idea. Studies combining experience sampling with measures from neuroimaging demonstrated increased activity in the DMN during off-task states (Christoff et al., 2009; Stawarczyk, Majerus, Maquet, & D’Argembeau, 2011). While others found tasks which mimic the off-task state, such as mental time travel and self-relevant or social cognition, modulate the DMN (for a review see Andrews-Hanna, Smallwood, & Spreng, 2014). Although these studies provide evidence of associations between the DMN and off-task thought, the broader view of this system as task-negative has been challenged (Spreng, 2012). For example, recent studies of connectivity during cognitive tasks have shown that the DMN communicates selectively with task positive systems (Krieger-Redwood et al., 2016), and that DMN activity can increase during cognitive task that require internally represented information or task rules (Crittenden, Mitchell, & Duncan, 2015; Murphy et al., 2018; 2019; Vatansever, Menon, & Stamatakis, 2017). If the functions of the DMN are not task-negative *per se* then this would undermine a simple one-to-one mapping of DMN activity and content processed in the off-task state.

An alternative perspective on the functional role of DMN in cognition emerges from a consideration of its status within the broader cortical landscape (Van Den Heuvel & Sporns, 2011). In humans and other primates, regions of the DMN are located at points on the cortex that are maximally distant from regions of unimodal cortex, in both functional and structural space (Margulies et al., 2016). Increased topographical distance from unimodal systems may help segregate neural signals in the DMN from sensory input, explaining why neural activity within this network has a unique relationship to task behaviour (i.e. task-induced deactivations). The cortical organisation giving rise to functional isolation is thought to represent a hierarchy which supports the progressive integration of information from unimodal regions to transmodal regions, the most extreme of which are in the DMN, supporting increasingly abstract neural representations (Mesulam, 1998). In this view, the DMN reflects a set of closely allied hubs that form an integrative cortical backbone vital for the maintenance of cortical dynamics (Van Den Heuvel & Sporns, 2011). If the DMN reflects a collection of integrative hubs, then it is likely to be linked to broadly distributed patterns of cortical activity that may determine relatively global features of ongoing cognition, rather than specific aspects of mental content.

In a prior study (Sormaz et al., 2018), we re-evaluated the relationship between neural activity and momentary patterns of ongoing thought, taking advantage of both recent advances in experience sampling - that describe dimensions underlying multiple aspects of self-reported experience - and imaging analysis methods. Neural function was recorded using functional magnetic resonance imaging (fMRI) while participants alternated between blocks of 0-back and 1-back working memory. During working maintenance (i.e. the 1-back condition), representational similarity analysis indicated that patterns of activity within the DMN were associated with the level of detail in experience reported by participants. Importantly, activity within the DMN was linked to relative levels of detail in experience during the maintenance of visual spatial information in working memory, and so contributed to participant’s experiences when they were more engaged with the task. These results therefore challenge notions of the DMN as contributing purely to task irrelevant cognition.

An alternative possibility is that the role of the DMN in ongoing thought emerges from contributions to specific features of off-task experiences, such as self-relevant or social content (Smallwood et al., 2016; Stawarczyk et al., 2011; Wong et al., 2017). This possibility is hard to rule out in our prior study since neural activity was sampled in a working memory paradigm, when the content of ongoing cognition is most dissimilar to those that emerge in the off-task state (Konishi, Brown, Battaglini, & Smallwood, 2017). Therefore, any relationship that was specific to particular aspects of content (such as it’s self-relevance) could have been obscured in our prior study because the visuo-spatial memory task lacked meaningful semantic or personally applicable content. To test this possibility, the current study examined whether the relationship between off-task experiences in the laboratory can be predicted by the neural responses that occur in the DMN when participants make trait judgements about themselves and a significant other. This manipulation has been shown to modulate activity within the DMN (Mitchell, Banaji, & Macrae, 2005), and can lead to increases in off-task thought in the lab (Stawarczyk, Cassol, & D’Argembeau, 2013). Notably, if DMN activity during off-task states reflects processes specific to content in that state, then the expression of self-relevant or social off-task mental content in the laboratory, should be linked to the recruitment of the DMN during trait adjective judgments. In contrast, if the DMN is important for broader aspects of ongoing experience, rather than just its contents, then neural activity within this system could show a positive correlation with descriptions of more abstract features of ongoing thought in the laboratory (i.e. its detail).

## 2. Methods

### 2.1. Participants

In total, 207 participants took part in a multidimensional experience sampling (MDES) study in the laboratory and the resting-state and structural scans (females = 132, M_age_ = 20.2, SD= 2.35 years). Out of these, 65 participants returned for an additional self-reference fMRI experiment (41 females; M_age_ = 21.4, SD = 2.67). The study received ethical approval from the York Neuroimaging Centre and the University of York Psychology Department.

### 2.2. Multidimensional experience sampling (MDES)

In the laboratory we used a block design to assess the contents of experience during a simple working n-back task (0-back; 1-back) (for prior published examples of this task see Konishi et al., 2015; Sormaz et al., 2018; Wang et al., 2018) across three separate sessions. Participants completed target and non-target trials. In non-target trials, a pair of shapes appeared on screen. Following an unpredictable sequence of non-target trials, a target trial was presented in which participants had to make a manual response. In the 0-back condition, the target was flanked by one of two shapes, and participants had to indicate which shape matched the target shape by pressing the appropriate button. In the 1-back condition, the target was flanked by two question marks, and participants had to respond depending on which side the target shape had been on during the prior trial. There were 8 blocks in one session and each block consisted of two to four mini-blocks. Each block contained either the 0-back or the 1-back condition. Each mini-block consisted of one target trial and a varied number of non-target trials preceding the target trial (between one and six non-target trials). Conditions were counterbalanced across participants. In total, the eight blocks lasted around 25 min.

In order to sample different features of participants’ ongoing experiences during the n-back task, we used multidimensional experience samples (MDES; Smallwood et al., 2016). This technique uses self-report data to assess the contents of experience on a number of dimensions. Participants answered 13 questions (see Table 1) on a 4-point scale ranging from 0-1. The first thought probe asked participants to rate their level of task focus (“My thoughts were focused on the task I was performing”) on a sliding scale from 0 (completely off-task) to 1 (completely on task). Participants then answered 12 randomly presented questions regarding the content and form of their experience in the moments just before they answered the first thought probe (on level of task focus). MDES probes occurred on a quasi-random basis to minimise the likelihood of anticipating the probes. At the moment of target presentation there was a 20% chance of a MDES thought probe being presented instead of a target, with a maximum of one probe per block of the 0-back/1-back task. In each of the three sessions, an average of 14.07 (SD = 3.30, range = 6–25) MDES probes occurred; in the 0-back condition, an average of 7.02 (SD = 2.36, range = 2–14) MDES probes occurred; and in the 1-back condition, an average of 7.04 (SD = 2.24, range = 1–15) occurred (average of 42 probes per participant across all three sessions).

**Table 1.**
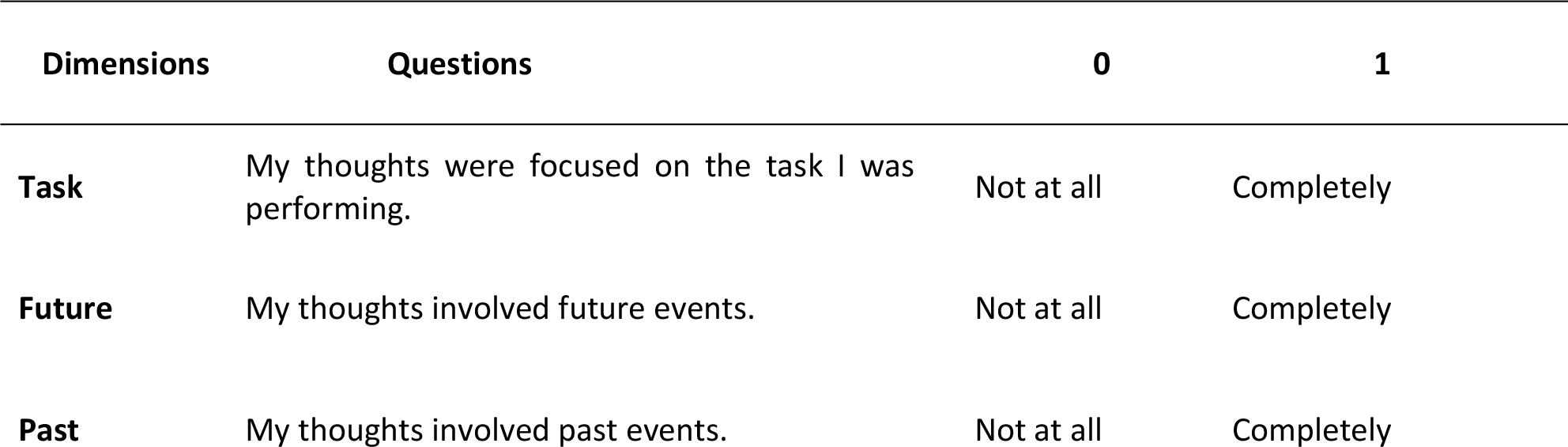
Multi-Dimensional Experience Sampling questions used this experiment.

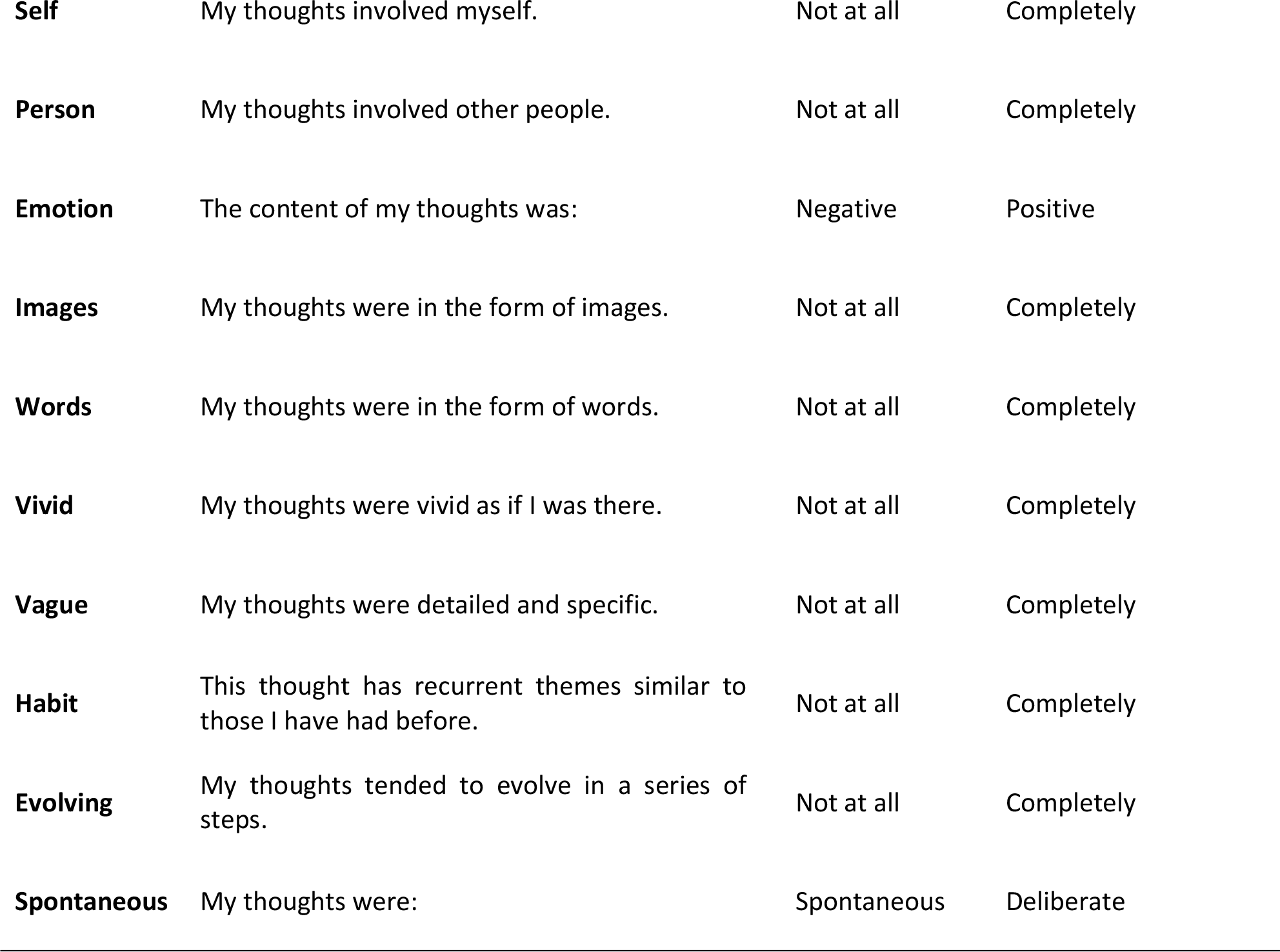

### 2.3. Resting-state fMRI

#### 2.3.1. Procedure

All 207 subjects completed a 9-minute functional connectivity MRI scan during which they were asked to rest in the scanner with their eyes open.

#### 2.3.2. Resting-state fMRI acquisition

As with the functional experiment, a Magnex head-dedicated gradient insert coil was used in conjunction with a birdcage, radio-frequency coil tuned to 127.4 MHz. For the resting-state data, a gradient-echo EPI sequence was used to collect data from 60 axial slices with an interleaved (bottom-up) acquisition order with the following parameters: TR = 3 s, TE = minimum full, volumes = 180, flip angle = 90°, matrix size = 64 × 64, FOV = 192 × 192 mm, voxel size = 3 × 3 × 3 mm. Functional images were co-registered onto a T1-weighted anatomical image from each participant (TR = 7.8 s, TE = 3 ms, FOV = 290 mm × 290 mm, matrix size = 256 mm × 256 mm, voxel size = 1 mm × 1 mm × 1 mm).

### 2.4. Self-reference fMRI task (n=65)

#### 2.4.1. Procedure

In the scanner, participants completed a trait-judgement paradigm where participants had to associate certain traits to themselves (‘Self’) or another person (‘Other’). Trials began with a 9 second fixation cross, followed by a 6 second presentation of a neutral valence adjective taken from a set of norms (Anderson, 1968). Participants decided whether the word applied to them (Self) or Barack Obama (Other). For instance, does ‘honest’ apply to Barack Obama? Responses (yes/no) were made after the six-second contemplating period using a button response. In both conditions 15 trials occurred and words were fully counterbalanced.

#### 2.4.2. MRI acquisition

Structural and functional data were acquired using a 3T GE HDx Excite MRI scanner with an 8-channel phased array head coil (GE) tuned to 127.4 MHz, at the York Neuroimaging Centre. Structural MRI acquisition was based on a T1-weighted 3D fast spoiled gradient echo sequence (TR = 7.8 s, TE = min full, flip angle= 20°, matrix size = 256 × 256, 176 slices, voxel size = 1.13 × 1.13 × 1 mm). Functional data was recorded using single-shot 2D gradient-echo-planar imaging (TR = 3 s, TE = minimum full, flip angle = 90°, matrix size = 64 × 64, 60 slices, voxel size = 3 × 3 × 3 mm^3^, 180 volumes). A FLAIR scan with the same orientation as the functional scans was collected to improve co-registration between scans.

### 2.5. Data Analysis

#### 2.5.1. Principal component analysis (PCA) of MDES data

We created a single 8694-by-13 matrix in which each row was a MDES probe (8694 = 207 participants × 42 probes) and each column was a question. We used PCA with varimax rotation in SPSS version 24 to reduce the dimensionality of these data using scree plots to determine that 4 principal components should be extracted (see Fig S1A). The resulting data were used as regressors of interest at the participant level in the self-reference experiment. The component names were decided by naming each after the highest loading item if the component was anchored at one end (e.g. Detail). If they were anchored by opposing scores (e.g., Words and Images) we named them based on the distinction that these loadings imply (e.g. Modality).

#### 2.5.2. Self-reference task analysis

For the analysis of neural activity during the self-reference task, functional and structural data were pre-processed and analysed using FMRIB’s Software Library (FSL version 5.0, http://fsl.fmrib.ox.ac.uk/fsl/fslwiki/FEAT/). Individual FLAIR and T1 weighted structural brain images were extracted using BET (Brain Extraction Tool). Structural images were registered to the MNI-152 template using FMRIB’s Linear Image Registration Tool (FLIRT). The functional data were pre-processed and analysed using the FMRI Expert Analysis Tool (FEAT). Individual subject analysis involved: motion correction using MCFLIRT; slice-timing correction using Fourier space time-series phase-shifting; spatial smoothing using a Gaussian kernel of FWHM 6mm; grand-mean intensity normalisation of the entire 4D dataset by a single multiplicative factor; highpass temporal filtering (Gaussian-weighted least-squares straight line fitting, with sigma = 100 s); Gaussian lowpass temporal filtering, with sigma = 2.8s.

First level analyses modelled the two experimental conditions (self and other). For each condition, these were modelled as six-second boxcar regressors modelling the entire presentation of the trait adjective (e.g., ‘honest’), during which participants had to think about whether that trait applied to them (Self condition) or Barack Obama (Other condition). The response to each condition was contrasted against rest. Boxcar regressors for each condition in the general linear model were convolved with a double gamma hemodynamic response function (FEAT, FSL). Regressors of no interest were also included to account for head motion. First level effects were entered into a group analysis using a mixed-effects design (FLAME, http://www.fmrib.ox.ac.uk/fsl). Z-stat maps were generated for each EV; Self-reference and Other-reference. These maps were then registered to a high resolution T1-anatomical image and then onto the standard MNI brain (ICBM152).

We performed two group level analyses on the task data using FLAME. To identify the spatial distribution of the regions modulated by the nature of the referent during trait adjective assessment, we compared periods of Self with Other (and the reverse). Both spatial maps are thresholded at Z = 3.1 and are controlled for multiple comparisons (p < .05, FWE). To understand the relationship between neural processing during each referent and experience in the laboratory, we performed group level regressions in which the spatial maps for each contrast where the dependent variables and the mean loadings for each individual for each MDES component were included as between participant continuous explanatory variables.

#### 2.5.3. Resting state fMRI analyses

Resting-state data pre-processing and statistical analyses were carried out using the SPM software package (Version 12.0), based on the MATLAB platform (Version 15a). For pre-processing, functional volumes were slice-time and motion-corrected, co-registered to the high-resolution structural image, spatially normalised to the Montreal Neurological Institute (MNI) space using the unified-segmentation algorithm (Ashburner & Friston, 2005), and smoothed with an 8 mm FWHM Gaussian kernel. With the goal of ensuring that motion and other artefacts did not confound our data, we first employed an extensive motion-correction procedure and denoising steps, comparable to those reported in the literature (Ciric et al., 2017). In addition to the removal of six realignment parameters and their second-order derivatives using the general linear model (GLM) (Friston et al., 1996), a linear de-trending term was applied as well as the CompCor method that removed five principle components of the signal from white matter and cerebrospinal fluid (Behzadi et al., 2007). Moreover, the volumes affected by motion were identified and scrubbed based on the conservative settings of motion greater than 0.5 mm and global signal changes larger than z = 3. Out of the 207 participants, a total of fifteen participants, who had more than 15% of their data affected by motion was excluded from the analysis (Power et al., 2014). Therefore, all resting-state analyses reported are based on a sample size of 192 participants. Finally, a band-pass filter between 0.009 Hz and 0.08 Hz was employed in order to focus on low frequency fluctuations.

Following this procedure, two seed regions were generated from the univariate maps of Self>Other and Other>Self regressed against our detail PCA (see Figure 2). This yielded two clusters centred on caudal posterior cingulate cortex for Self>Other [MNI: −2, −58, 26] and intracalcarine cortex for Other>Self [MNI 16, −84, 4]. The Conn functional connectivity toolbox (Version 15.h) (Whitfield-Gabrieli & Neston-Castanon, 2012) was used to perform seed-based functional connectivity analyses for each subject using the average signal from the spheres placed on the MNI coordinates for the two regions of interest (ROIs) described above. All reported clusters were corrected for multiple comparisons using the Family-Wise Error (FWE) detection technique at the .05 level of significance (uncorrected at the voxel-level, .001 level of significance).

## 3. Results

We applied PCA to the trial level data produced by MDES. This method describes the underlying dimensions within the patterns of responses in the experience sampling data. These can be broadly understood to be ‘patterns of thought’ captured in our study. Based on the scree plot (see Figure S1), we identified four components that described 54.94% of the variance in our data and the loading of each item in each dimension are presented in the form of word clouds in the grey box in Figure 1. Component 1 described high loadings on detailed and evolving thoughts (Detail). This corresponds to the pattern of experience linked to the DMN in our prior study (Sormaz et al., 2018). Component 2 described low loadings on the task and deliberate thought and higher loadings on self, future and person thoughts (Task). This component corresponds to the off-task state since it places personal and socially relevant content in opposition to a deliberate focus on the task. Component 3 described positive loadings on images and negative loadings on words and so corresponded to variation in the modality of experience (Modality). Component 4 describes high loadings on positive affect (Emotion). Note that these patterns are derived from the application of MDES to the larger sample of 207 individuals as this provides the most stable estimation of these dimensions. We repeated this analysis pipeline on the subset of individuals who also performed the self-reference task, yielding very similar scree plot (see S1B) and pattern of loadings (see Figure S2B). In addition, in our prior study we demonstrated that the patterns derived using PCA generalised to different data collected in the same individuals (see Sormaz and colleagues for further details).

**Figure 1.**
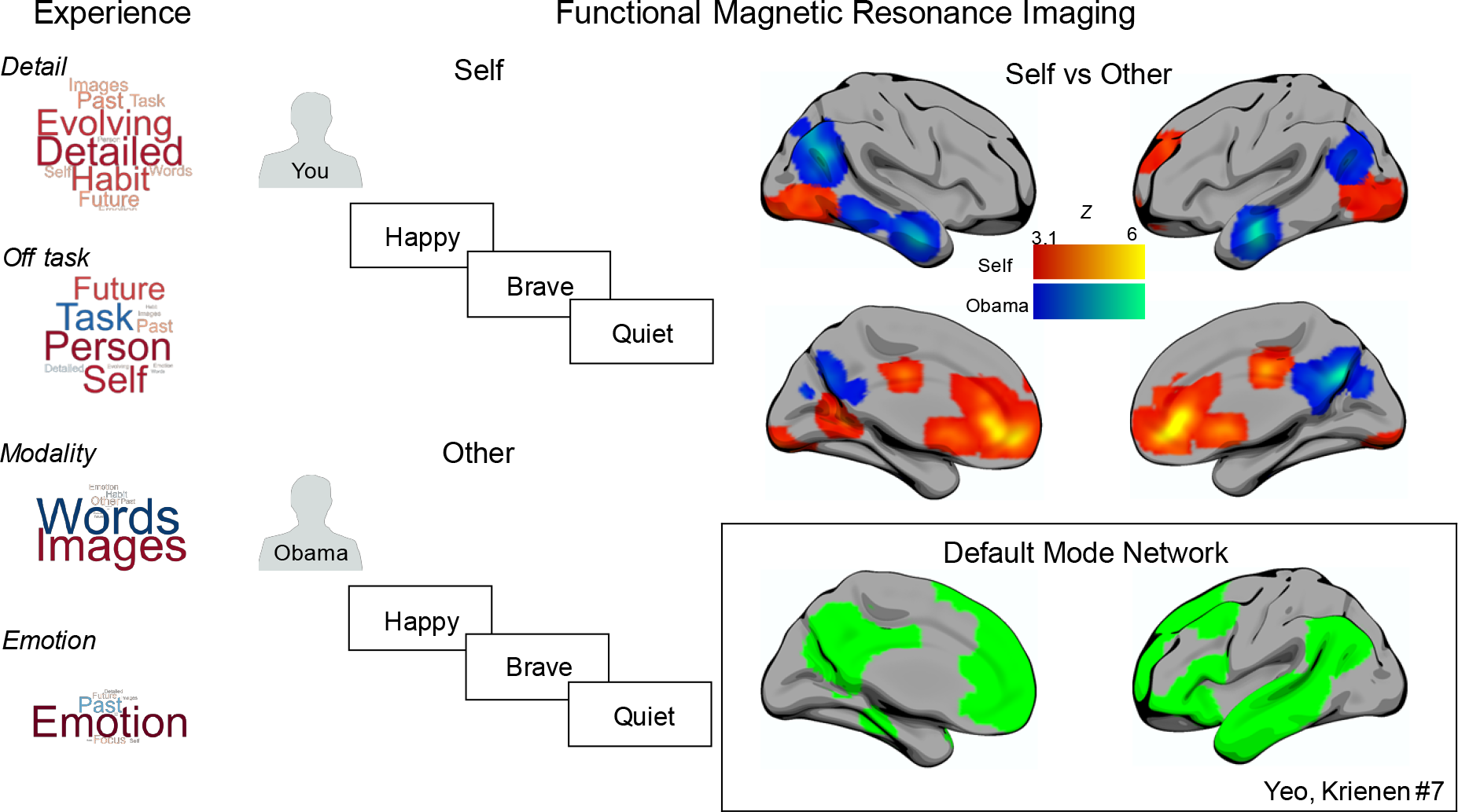
Patterns of thought during the laboratory and the modulation of neural activity during the self-reference task. The panel on the left-hand side displays the components of thought determined by the application of Principal Components Analysis to the Multi-Dimensional Experience Sampling (MDES) using a sample of 207 individuals. The size of the word describes the strength of the loading and the colour identifies whether the loadings is positive (warm) or negative (cool). The central panel describes the block design used to assess neural processing during a self-reference task. The brains display the spatial distribution of the regions identified through a contrast of activity between self and other referent conditions. Warm colours highlight regions showing more activity during Self judgements and cool colours show regions showing more activity in Other judgements. The spatial maps are thresholded at Z = 3.1 and are controlled for multiple comparisons (p < .05, FWE). The box in the lower right hand corner shows the DMN as defined in the 7-network solution by Yeo, Krienen et al (2011).

The comparison of neural activity when participants made self-reference judgements in the scanner is presented in the right hand panel of Figure 1. Manipulation of the referent (Self or Other) upon which trait judgements were made successfully modulates activity within the DMN. When making decisions about the Self, activity was higher in regions of ventromedial prefrontal cortex, as well as in two regions of posterior cingulate cortex. When relating adjectives to Barak Obama, activity was higher in regions of angular gyrus, lateral temporal cortex, as well as in the core of the posterior cingulate. All the regions modulated by the referent of the trait adjective judgement, with the exception of ventral lateral occipital activity, fall within the DMN as defined by Yeo and colleagues (see inset to Figure 1). The modulation of activity within the DMN by different traits is broadly consistent with the general importance for the temporal lobe regions in social cognition and from prior studies similar to this one (Andrews-Hannah et al., 2014). Likewise meta-analyses suggest greater involvement of frontal DMN regions during self-relevant processing (Hu et al., 2016).

Next, we considered how patterns of experience in the lab are related to the neural activity during the self-reference task. We performed separate group level regressions in which the dependent variables were maps of neural activity from each condition (Self and Other). Each individual’s average score on each MDES component from the laboratory was included as between individual explanatory variables. This analysis identified neural regions in both task conditions that increased activity in proportion to how detailed an individual’s thought was in the lab. These are presented in Figure 2. In the Other condition, participants whose thoughts were more detailed had greater activity in a region of medial occipital cortex. This is presented in cool colours. In addition, during the Self condition, recruitment of caudal posterior cingulate cortex (PCC) - a core hub of the DMN - was greater for participants whose laboratory experiences were characterised as detailed and evolving. This is presented in warm colours in Figure 2. Scatter plots describing both effects are presented in the left-hand lower panel.

**Figure 2.**
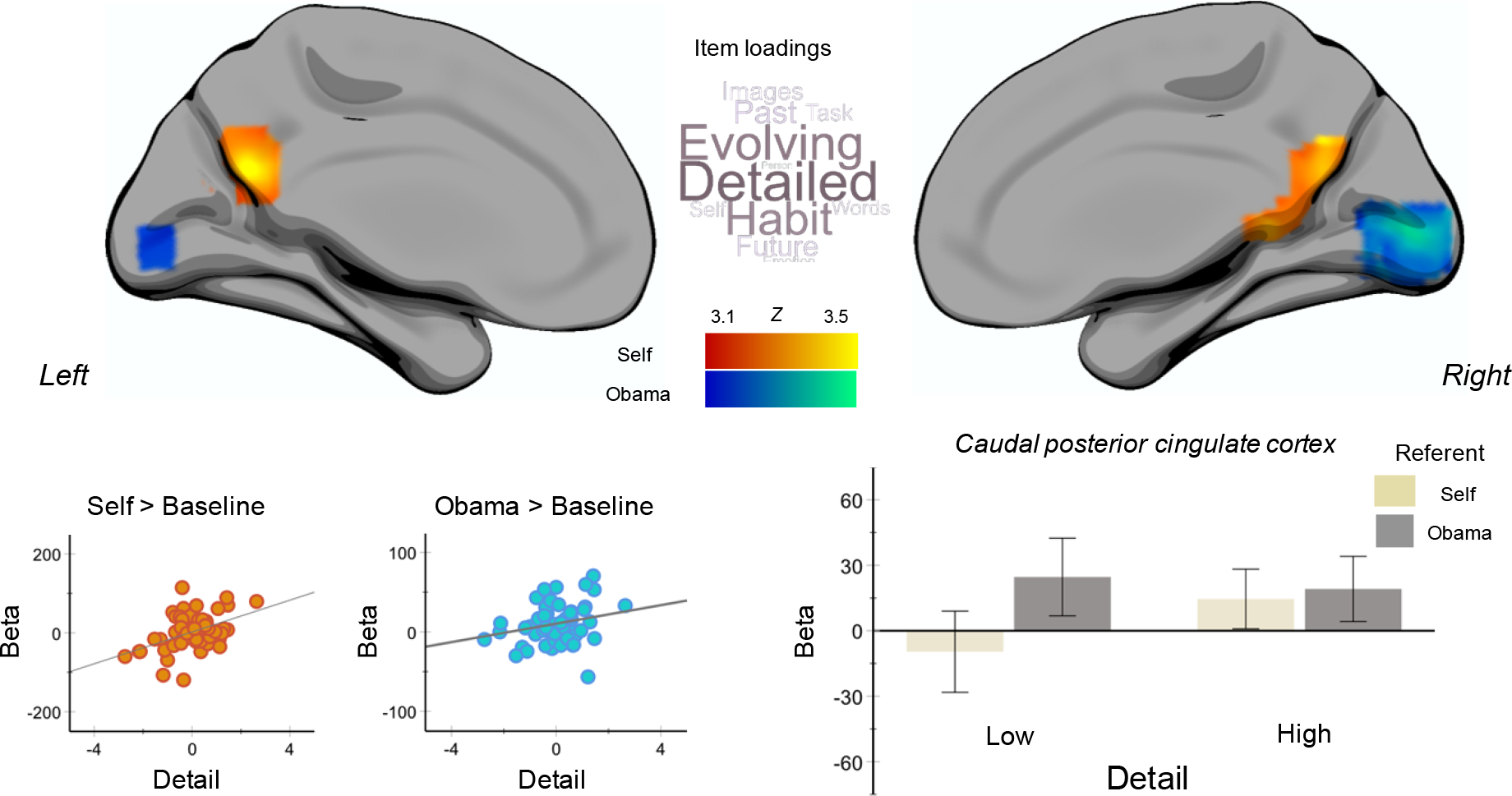
The relationship between patterns of ongoing thought in the laboratory and neural activity during trait adjective judgements. Warm colours describe regions whose neural activity during self-trait judgements was correlated with levels of detail in ongoing experience in the lab. Cool colours describe regions whose neural activity during other-trait judgements was correlated with the level of detail in ongoing thought. Spatial maps were thresholded at Z = 3.1 controlling for multiple comparisons p < .05 (FWE). The scatter plots display the association between individual’s detail component score and neural activity extracted from posterior cingulate cortex (PCC) (warm colours) and medial occipital cortex (cool colours) respectively. Notably, the PCC cluster also showed a selective increase when making trait judgements regarding Barak Obama relative to the Self (see Figure 1 and Figure S3). The bar graph in the grey inset shows the signal in the rostral PCC cluster in each condition (Self / Other) plotted for individuals whose thoughts in the lab were below (Low) and above (High) the median for the Detail dimension. Thus, individuals who engaged in detailed thought within the laboratory recruited caudal posterior cingulate cortex above baseline when making trait adjective judgements based on either Self or Barack Obama as the source of the judgement. The error bars indicate the 95% C.I. The word cloud summarises the pattern of loadings that correspond to the detail component. Larger fonts describe a higher loading.

Notably, the PCC region of the DMN that showed greater activity during self-reference when participants experiences were detailed and evolving (Figure 2), also tended to show more activation to rating adjectives of another (Obama) rather than the self (see Figure 1 and the results of a formal conjunction analysis in Supplementary Figure S4). This indicates that the region that shows an association with detail in the lab during self-reference tends to have higher activity when considering whether adjectives applied to Obama. To understand the relationship between these two spatially overlapping effects, we plotted the signal change in this region of caudal posterior cingulate cortex in both conditions, separating individuals into those who were above (High) and below (Low) the median for detail thinking in the lab (see lower right-hand panel in Figure 3). While individuals low on detailed thinking only showed greater activity in caudal posterior cingulate when rating adjectives with respect to Obama (Low group: PCA1 and Other (r = - .21, *p* = .15), PCA1 and Self (r = .346, *p* = .033), Other and Self (r = .298, *p* = .058)), for individuals high on detail the activity in this region was above baseline in both conditions (High group: PCA1 and Other (r = .231, *p* = .12), PCA1 and Self (r = .34, *p* = .037), Other and Self (r = .453, *p* = .008)).

**Figure 3.**
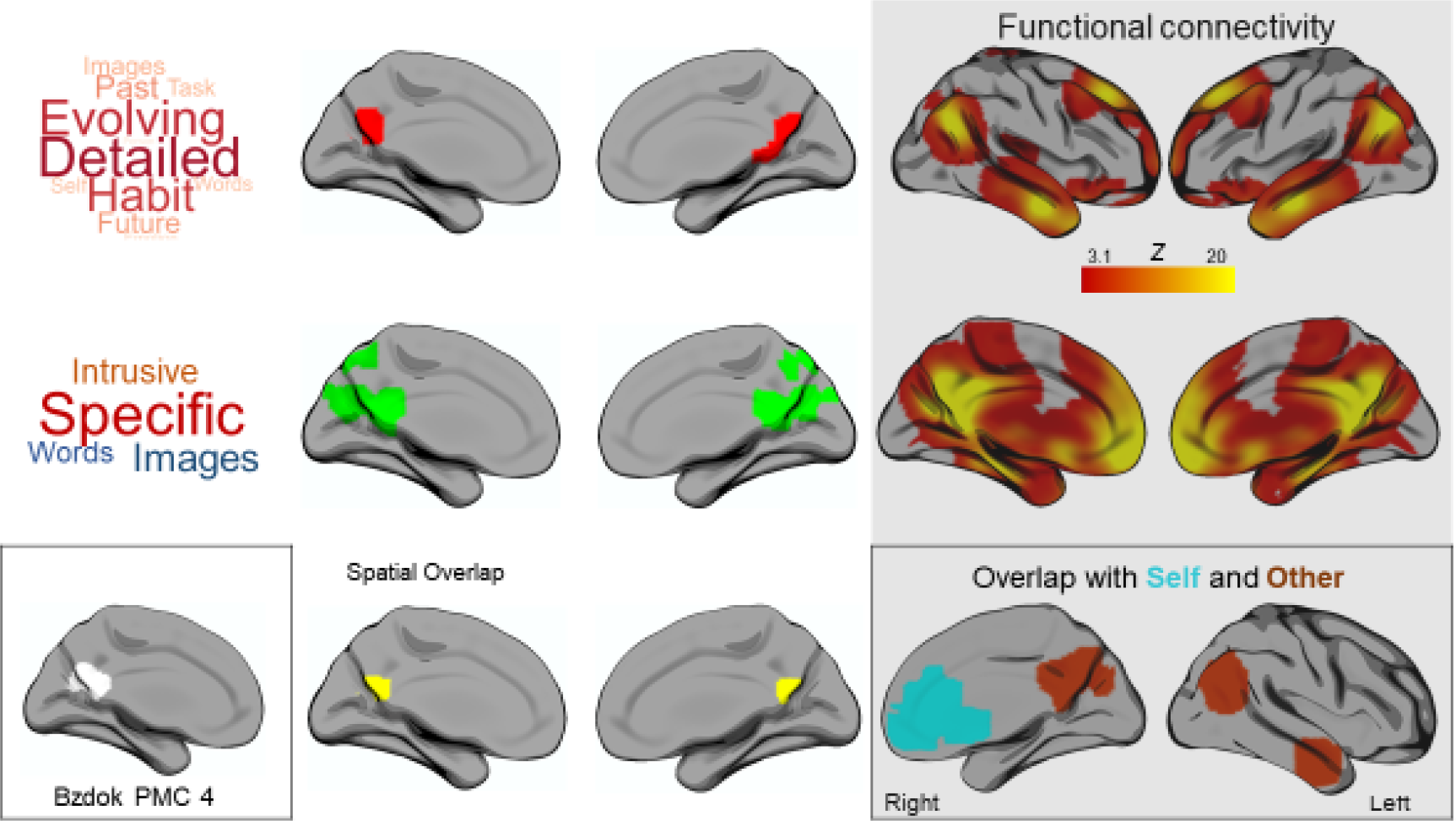
Converging evidence for a relationship between caudal posterior cingulate cortex and levels of specific detail in ongoing experience. The left-hand panel shows the spatial distribution of areas associated with patterns of detailed thought in the lab and during self-reference in the scanner (red), the results of an individual difference analysis from our prior study (Smallwood et al., 2016) showing associations between medial temporal lobe connectivity and the posterior cingulate indicating more detailed thoughts (green). These overlap in the area in yellow indicated in the lower panel. The inset shows a region of posterior cingulate cortex that was identified using decompositions based on both task and task-free data which was related to a combination of mental processes including language processing (Bzdok et al., 2015). The left-hand panel shows the functional connectivity of this region (higher correlations indicated in warmer colour). This pattern of connectivity captures the regions that show a pattern of modulation by both Self and Other referents during the self-reference task as can be seen in the lower left inset. The functional connectivity maps are t corrected for multiple comparisons using the Family-Wise Error (FWE) detection technique at the .05 level of significance (uncorrected at the voxel-level, .001 level of significance)

In contrast, to the effects of detail there were no regions in a whole brain analysis significantly associated with the extent of off-task thinking in the lab, and post hoc analysis indicated no association between neural processing in either the posterior cingulate cortex, or the lingual gyrus (all r-values < .15, all p-values > .6). Thus, while no association was observed between patterns of off-task content in the laboratory, we found evidence that signals in the DMN were related to individual differences in the tendency to increase activity within caudal posterior cortex during patterns of both self and other thinking.

While the method of whole brain analysis used in our prior analysis is well suited to the identification of positive relationships between patterns of neural activity and ongoing thought, it is possible that it may obscure important sub-threshold relationships between off-task thinking and patterns of neural activity. To rule out the possibility that our failure to find evidence of associations with off-task thinking is a Type 2 error we conducted a regions of interest (ROI) analysis targeting all cortical areas that showed the ability to dissociate self-relevant and social cognition. This was a total of 10 ROIs and we compared their signal change (relative to baseline) in both conditions of the experiment to both patterns of detail and off-task thinking (a total of 20 comparisons). This analysis revealed significant positive associations with detailed thinking in the posterior cingulate cortex in two regions linked to social cognition (posterior cingulate cortex, r = .31, p <.05, and left angular gyrus, r = .26, p =.051, as well as in the lateral occipital cortex during self-relevance (r = .31, p<.05). No regions showed a significant negative correlation. In contrast, only one region was the neural signal linked to off-task thought, with a region of anterior medial prefrontal cortex showing a significant negative association with neural processing during self-relevance (r = - .31, p<.05). Finally, a Bayesian analysis revealed that the evidence in support of the null hypothesis was greater than 3, moderate evidence for the null hypothesis, for 17 out of the 20 comparisons with off-task thought. Table 2 presented the distribution of r-values and Bayes factors for each of the correlations.

**Table 2.**
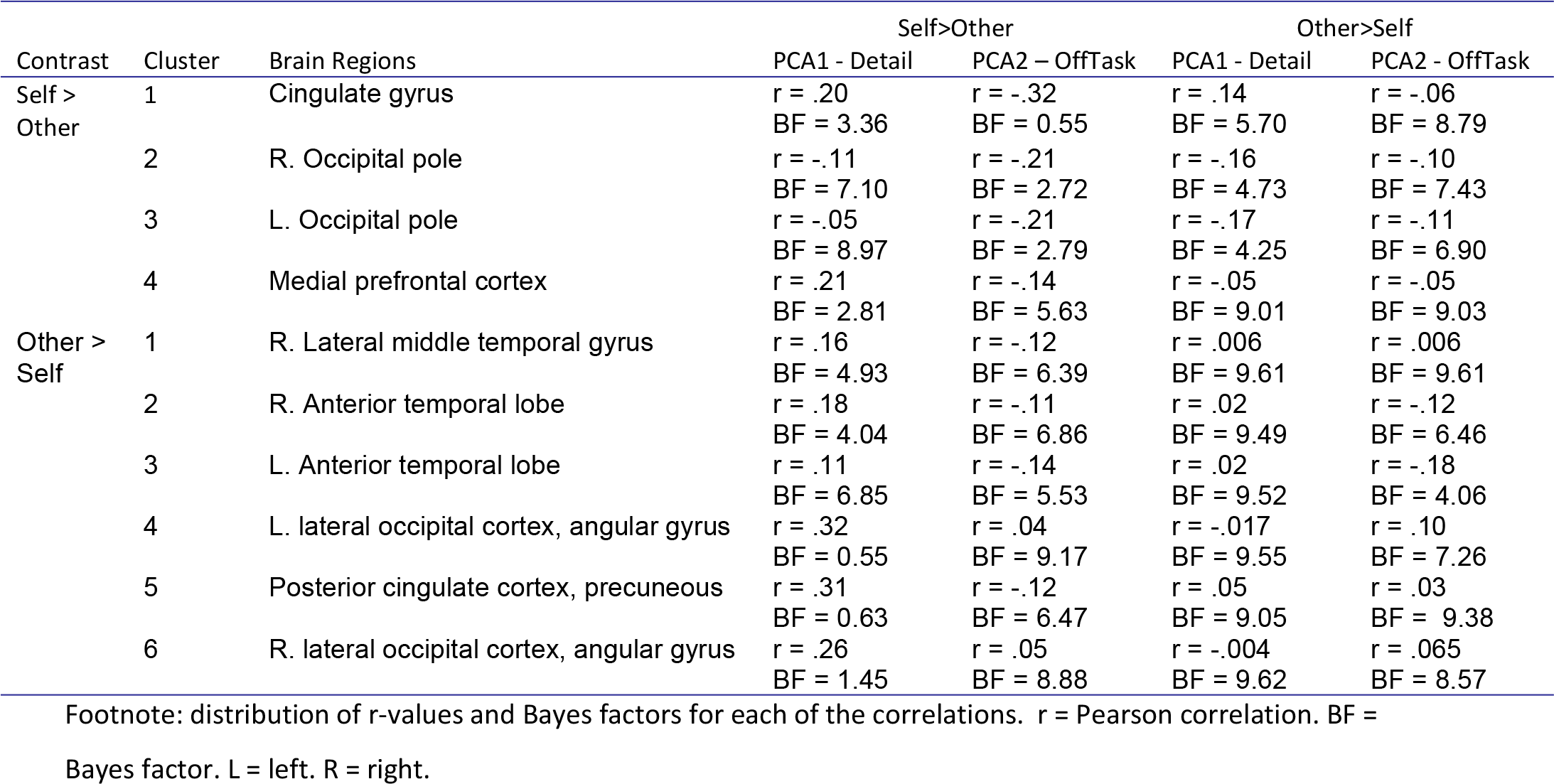
Bayesian analysis results

| Contrast | Cluster | Brain Regions | Self>Other |  | Other>Self |  |
| --- | --- | --- | --- | --- | --- | --- |
|  |  |  | PCA1 - Detail | PCA2 – OffTask | PCA1 - Detail | PCA2 - OffTask |
| Self > Other | 1 | Cingulate gyrus | r = .20<br>BF = 3.36 | r = -.32<br>BF = 0.55 | r = .14<br>BF = 5.70 | r = -.06<br>BF = 8.79 |
|  | 2 | R. Occipital pole | r = -.11<br>BF = 7.10 | r = -.21<br>BF = 2.72 | r = -.16<br>BF = 4.73 | r = -.10<br>BF = 7.43 |
|  | 3 | L. Occipital pole | r = -.05<br>BF = 8.97 | r = -.21<br>BF = 2.79 | r = -.17<br>BF = 4.25 | r = -.11<br>BF = 6.90 |
|  | 4 | Medial prefrontal cortex | r = .21<br>BF = 2.81 | r = -.14<br>BF = 5.63 | r = -.05<br>BF = 9.01 | r = -.05<br>BF = 9.03 |
| Other > Self | 1 | R. Lateral middle temporal gyrus | r = .16<br>BF = 4.93 | r = -.12<br>BF = 6.39 | r = .006<br>BF = 9.61 | r = .006<br>BF = 9.61 |
|  | 2 | R. Anterior temporal lobe | r = .18<br>BF = 4.04 | r = -.11<br>BF = 6.86 | r = .02<br>BF = 9.49 | r = -.12<br>BF = 6.46 |
|  | 3 | L. Anterior temporal lobe | r = .11<br>BF = 6.85 | r = -.14<br>BF = 5.53 | r = .02<br>BF = 9.52 | r = -.18<br>BF = 4.06 |
|  | 4 | L. lateral occipital cortex, angular gyrus | r = .32<br>BF = 0.55 | r = .04<br>BF = 9.17 | r = -.017<br>BF = 9.55 | r = .10<br>BF = 7.26 |
|  | 5 | Posterior cingulate cortex, precuneus | r = .31<br>BF = 0.63 | r = -.12<br>BF = 6.47 | r = .05<br>BF = 9.05 | r = .03<br>BF = 9.38 |
|  | 6 | R. lateral occipital cortex, angular gyrus | r = .26<br>BF = 1.45 | r = .05<br>BF = 8.88 | r = -.004<br>BF = 9.62 | r = .065<br>BF = 8.57 |

Footnote: distribution of r-values and Bayes factors for each of the correlations. r = Pearson correlation. BF = Bayes factor. L = left. R = right.

Finally, to examine the generalisability of the association found between neural processing in caudal posterior cortex and detailed thought in the laboratory we compared the current data to a similar region identified in a prior study (Smallwood et al., 2016). This study analysed the resting-state connectivity of DMN nodes including those in the temporal lobe (the temporal pole and hippocampus) and their relationship to patterns of ongoing thought in the laboratory, using a similar laboratory paradigm to the one in the current experiment. Connectivity between a region of medial temporal lobe, and, a region of posterior cingulate cortex was stronger for individuals who had more specific patterns of ongoing thought. These spatial maps both overlapped within the caudal posterior cingulate (shown in yellow in the inset of Figure 3). This correspondence between neural evidence from the domain of connectivity and task-based fMRI provides converging evidence of a link between this posterior node in the DMN with patterns of ongoing thought that are detailed and specific. To place this region of caudal posterior cingulate in a broader cortical context, we examined its intrinsic connectivity architecture using resting-state data collected in the larger sample. Functional connectivity analyses revealed a pattern of correlation with many of the structures in the DMN including the posterior cingulate cortex, ventromedial prefrontal cortex, medial and lateral temporal cortex and angular gyrus (see the grey panel in Figure 3). Comparison of this pattern of functional connectivity with maps of regions modulated by the nature of the referent during trait judgements (Self > Other and the reverse) revealed that all but one region (lateral occipital cortex) fell within the connectivity map of this region. This was confirmed by a formal conjunction of the three maps (see the right hand inset).

In sum, our study found no association between the neural processing during Self or Other focused mentation and patterns of similar themed ongoing thought in the laboratory. Neural engagement during Self and Other trait processing, therefore, was not correlated to reports of off-task content, despite PCA highlighting these topics as characteristic themes of this state. Instead activity in a caudal aspect of the posterior cingulate during self-reference was linked to levels of detail in experiences in the lab. The absence of an association with off-task thought cannot be easily attributed to a lack of power in our study, since our Bayesian analysis suggests that the evidence in favour of the null hypothesis is reasonably strong. Likewise they cannot be discounted by virtue of a lack of sensitivity of the task focus component produced through our method of analysis, since broadly equivalent patterns of experience, generated by the same method, vary with objectives measures including neural (Sormaz et al., 2018) and pupilometric (Konishi et al., 2017) data.

## 4. Discussion

While our data are inconsistent with a simple role for the DMN in self-relevant content of off-task thought, they do not exclude a role in broader features of cognition, such as its detail or meaning. This account draws support from studies probing the mechanics of cognition in tasks, especially those in the domain of memory. For example, patterns of activity within the posterior core of the DMN describe the vividness of episodic retrieval (Richter, Cooper, Bays, & Simons, 2016). They also code longer, more semantically relevant sections of narratives during television and audiobooks (Baldassanno et al., 2017). Neuro-stimulation is difficult in medial regions of the DMN, however, studies show that disrupting function in lateral aspects of the DMN within the angular gyrus impair source monitoring and confidence of episodic memory (Sestieri et al., 2013), and, in the semantic domain, impairs specific retrieval but not more difficult patterns of retrieval (Davey et al., 2015).

A role of the DMN in supporting broad features of cognition, such as the experienced level of detail, is consistent with neuroanatomical evidence situating this system as an integrative core located at the top of a neurocognitive hierarchy (Margulies et al., 2016). This topographical organisation is assumed to allow transmodal regions to be less tethered to specific modalities of input and so better able to integrate from a broader range of cortical inputs (Buckner & Krienen, 2013). Accordingly the capacity of regions such as the posterior cingulate to draw on neural signals from a wide range of cortical regions could enable it to support more elaborate and detailed mental representations (Ramanan et al., 2018), a process that is often described as scene construction (Hassabis & Maguire, 2007). These converging lines of evidence support a role for the DMN in ongoing thought that is related to broader features of cognition, such as how it is being represented, rather than more local features such as what it is about (Gonzalez-Garcia et al., 2018; Smallwood et al., 2016).

Prior observations of DMN activity within the off-task state (Christoff, Gordon, Smallwood, Smith, & Schooler, 2009; Stawarczyk et al., 2011), may, therefore have been at least partially mischaracterised. Rather than reflecting the act of processing particular content, the recruitment of structures within the DMN may describe patterns of ongoing thought with particular features, such as whether they have vivid, absorbing details. These mischaracterisations may have emerged for methodological reasons. Prior studies often have employed tasks with limited semantic or self-relevant content, and often with few cognitive demands, to measure ongoing thought. Under these conditions, being on-task may be boring and dull, while when we are off-task we can generate imaginative experiences that are richer in certain aspects. We overcome this limitation in the current study by sampling neural function in a relatively engaging self / trait adjective task and associating this with self-reports of experience in the lab. This allows neural processing to be described in a context when participants perform specific tasks and so allows for a more mechanistic constraint on interpretations of the functions of neural processing during unconstrained states. We also measured multiple features of ongoing thought, while studies typically focus on only a few aspects of experience, often using a series of forced binary choices (e.g. on vs. off-task). Although there are clear methodological advantages to simple experience sampling regimes, such as efficiency, they are less well suited to probing specific nuances of the complex landscape of ongoing thought. Our approach overcomes this problem by measuring experience along multiple dimensions and examining the latent structure contained by these data.

However, complex experience sampling with multiple items has weaknesses (Seli et al., 2018) - it is time consuming, and may lead to concerns of lack of validity given its intrusiveness or reactivity (Schooler, 2002). Fortunately some of these issues can be addressed by measurements made across multiple studies. For example, in the current study we acquired neural activity during self and other judgements and related these to experience measures recorded in the lab, while in our prior experiment ongoing neural activity was related to online measures of experience sampling (Sormaz et al., 2018). As both studies implicated the DMN in patterns of detailed ongoing thought, we can be confident that this conclusion does not rest on a factor unique to only one method – such as momentary changes in neural function occurring because of the momentary load induced by experience sampling (Schooler, 2002). Moving forward, it seems likely that accurately describing associations between neural function and ongoing thought, requires the characterisation of multiple dimensions of experience and the development of the methodological tools that enable this heterogeneous landscape to be explored (Christoff et al., 2016; Smallwood & Schooler, 2015). It is also important to consider whether our data has sufficient power to detect the effects that it did. Based on our sample size N=65, we have an 80% power to detect correlations in the .34 range and above (Hulley et al., 2013). The correlations identified in our ROI analyses were slightly below this level (in the range of .26-.31). Although our sample of 65 is relatively high for a task based fMRI experiment, future studies may wish to consider a larger sample size to ensure the robustness of the effects.

## Acknowledgments

This study was supported by awards to JS from the ERC (WANDERINGMINDS - 646927), the Volkswagen Foundation (Wandering Minds - 89440 and 89439) and the John Templeton Foundation, “Prospective Psychology Stage 2: A Research Competition”. EJ was supported by grant FLEXSEM-771863 from the ERC.

---

**Fig S1.**
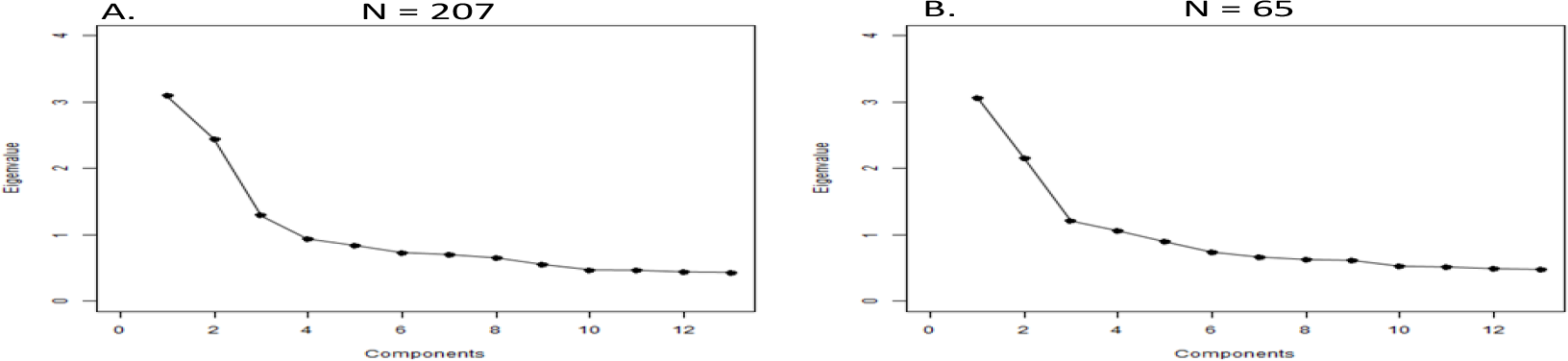
Scree plot describing the decomposition of experience sampling data in the lab for all 207 participants (A) and for a subset of these (N=65) who took part in the self-reference task (B). In both decompositions the first four components revealed a significant positive correlation (PCA1 r = .69, p = .009; PCA2 r = .72, p = .005; PCA3 r = .94, p < .001; PCA4 r = − .65, p =.02).

**Fig S2.**
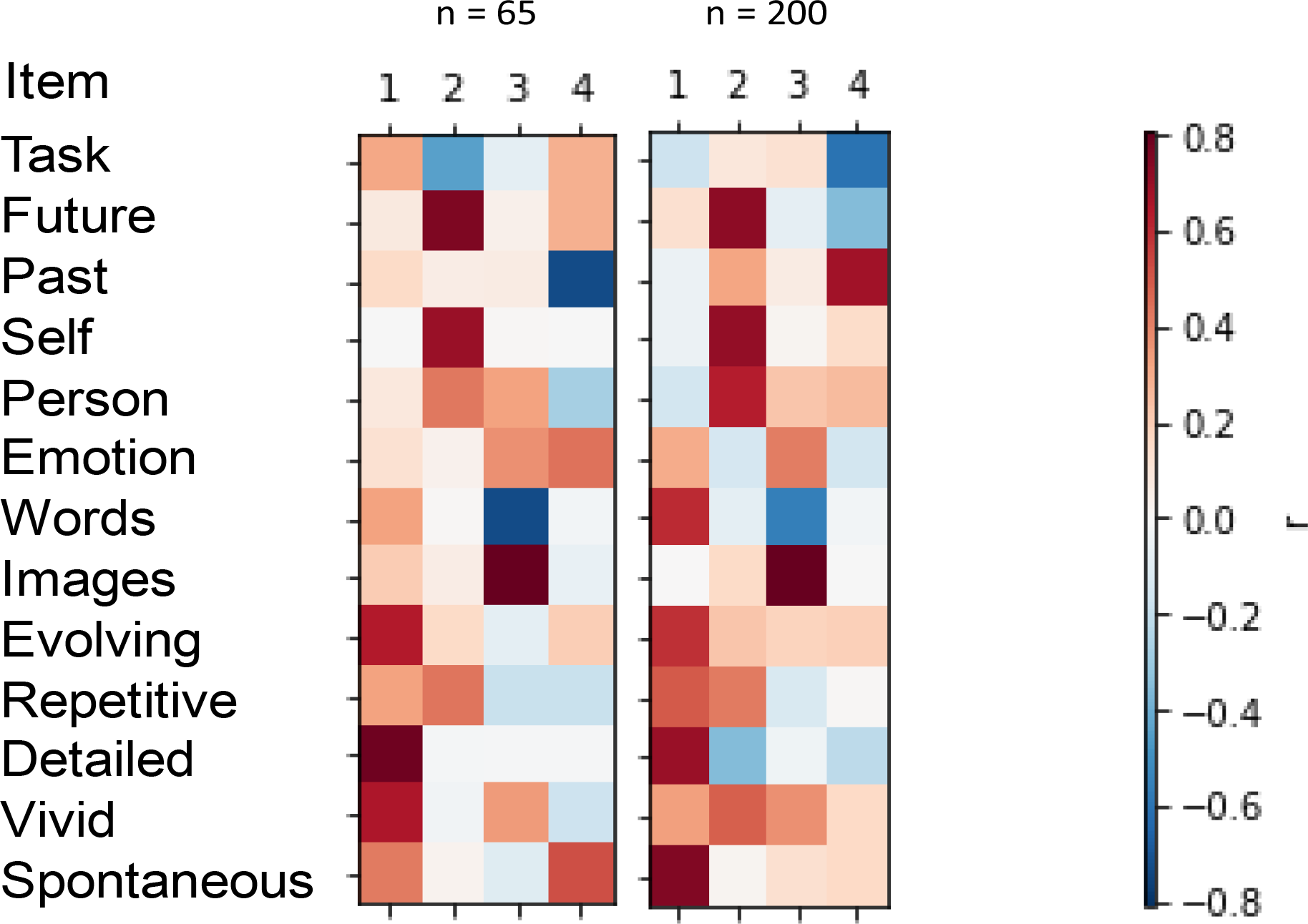
Heat maps describing the decomposition of experience sampling data in the lab for all 207 participants (left) and for a subset of these (N=65) who took part in the self-reference task (right). Correlations between the maps revealed a significant relationship between each component across the two datasets (PCA1 r = .69, p = .009; PCA2 r = .72, p = .005; PCA3 r = .94, p < .001; PCA4 r = −.65, p =.02).

**Figure S3.**
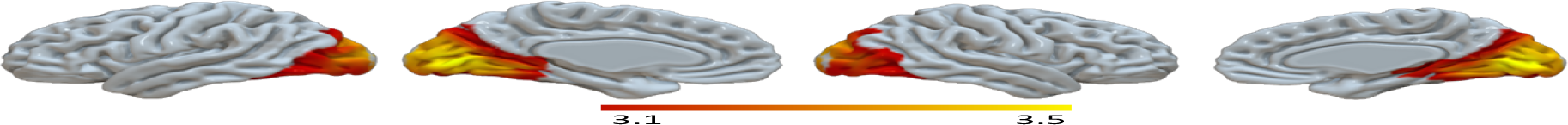
Functional connectivity of intracalcarine seed taken from Other>Rest regressed against detail component (figure 2). All reported clusters were corrected for multiple comparisons using the Family-Wise Error (FWE) detection technique at the .05 level of significance (uncorrected at the voxel-level, .001 level of significance).

**Figure S4.**
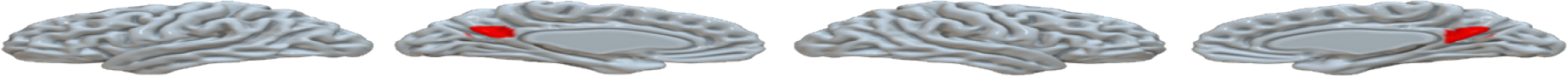
Conjunction of Other>Self whole-brain map (cool colour; figure 1) and Self>Rest regressed against Detail component (warm colours; figure 2) reveals cluster in caudal posterior cingulate cortex.

